# The impact of *Staphylococcus aureus* cell wall glycosylation on langerin recognition and Langerhans cell activation

**DOI:** 10.1101/2020.11.06.371559

**Authors:** A Hendriks, R van Dalen, S Ali, D Gerlach, GA van der Marel, FF Fuchsberger, P Aerts, CJC de Haas, A Peschel, C Rademacher, JAG van Strijp, JDC Codée, NM van Sorge

## Abstract

*Staphylococcus aureus* is the leading cause of skin and soft tissue infections. It remains incompletely understood how skin-resident immune cells respond to *S. aureus* invasion and contribute to an effective immune response. Langerhans cells (LCs), the only professional antigen-presenting cell type in the epidermis, sense *S. aureus* through their pattern-recognition receptor langerin, triggering a pro-inflammatory response. Langerin specifically recognizes the β-1,4-linked *N*-acetylglucosamine (β-GlcNAc) modification, which requires the glycosyltransferase TarS, on the cell wall glycopolymer Wall Teichoic Acid (WTA). Recently, an alternative WTA glycosyltransferase, TarP, was identified in methicillin-resistant *S. aureus* strains belonging to clonal complexes (CC) 5 and CC398. TarP also modifies WTA with β-GlcNAc but at the C-3 position of the WTA ribitol phosphate (RboP) subunit. Here, we aimed to unravel the impact of β-GlcNAc linkage position for langerin binding and LC activation. In addition, we performed structure-binding studies using a small panel of unique chemically-synthesized WTA molecules to assess langerin-WTA binding requirements. Using FITC-labeled recombinant human langerin and genetically-modified *S. aureus* strains, we observed that langerin similarly recognized bacteria that produce either TarS- or TarP-modified WTA. Furthermore, using chemically-synthesized WTA, representative of the different *S. aureus* WTA glycosylation patterns, established that β-GlcNAc is sufficient to confer langerin binding. Functionally, *tarP*-expressing *S. aureus* induce increased cytokine production and maturation of *in vitro*-generated LCs compared to *tarS*expressing *S. aureus*. Overall, our data suggest that LCs are able to sense all β-GlcNAc-WTA producing *S. aureus* strains, likely performing an important role as first responders upon *S. aureus* skin invasion.

## Introduction

*Staphylococcus aureus* is a Gram-positive bacterium that transiently colonizes an estimated 20% of the human population at different sites of the body, including the nasopharynx, skin and gastrointestinal tract. The skin is a common entry site for *S. aureus*, making it the leading cause of skin and soft tissue infections (SSTIs)(1). Consequently, efficient and rapid recognition of invading *S. aureus* by resident skin immune cells is critical for local eradication. When local immune defense fails, bacteria can disseminate into deeper tissues or even cause systemic infections, which are associated with high overall disease burden and mortality. The high recurrence of *S. aureus* SSTIs indicates that protective immune memory is absent, the reasons for which remain unknown. Indeed, there are no clear correlates of protection known for *S. aureus*, which has been a challenging aspect for vaccine development (2). A complete understanding of the local skin immune response to *S. aureus* may identify the factors that protect the host from (re-)infection, thereby providing critical insight for the development of a future *S. aureus* vaccine.

The skin contains a large arsenal of immune cells, which reside in different compartments within the skin. Langerhans cells (LCs), a highly specialized macrophage subset with dendritic cell-like functions, is the main antigen-presenting cell within the epidermis (3). Human LCs appear to have an important dual role in maintaining skin homeostasis, by balancing both tolerogenic responses towards skin commensals as well as pro-inflammatory responses to invading pathogens (4-9). However, the ability of LCs to recognize and respond to invading bacteria remains elusive due to their restricted expression of Toll-like receptors (10,11). C-type lectin receptors (CLRs) constitute another family of pattern-recognition receptors (PRRs), which are dedicated to the recognition of glycans (12). A signature CLR of LCs is langerin (CD207) (13). Langerin is a trimeric type II transmembrane receptor with specificity for sulfated and mannosylated glycans as well as β-glucans, which are recognized in a calcium-dependent manner (14-16). The direct downstream effects of receptor activation remain to be elucidated, since langerin only contains a short cytoplasmic tail without classical signaling motifs (13). It is generally assumed that langerin-bound cargo is endocytosed and processed for antigen presentation to CD4 T cells via major histocompatibility complex class II (MHC-II) (17-19).

Recent work demonstrated that langerin allows human LCs to discriminate *S. aureus* from other staphylococci through a specific interaction with glycosylated Wall Teichoic Acid (WTA) (20). WTA is a major component of the Gram-positive bacterial cell wall and a well-known immunogenic antigen for antibodies targeting *S. aureus* (21-23). *S. aureus* WTA consists of a polymerized ribitol phosphate (RboP) backbone that can be co-decorated with positively-charged D-alanine and *N*-acetylglucosamine (GlcNAc) residues. D-alanylation of WTA is highly regulated and impacts bacterial surface charge, thereby providing protection from host cationic antimicrobial peptides (AMPs) and the lipopeptide antibiotic daptomycin(24-27). WTA glycosylation can be mediated by different glycosyltransferases, resulting in distinct WTA glycoforms. Three different WTA glycoforms have been identified in *S. aureus*, which differ in the configuration and position of GlcNAc linkage. Langerin binding to *S. aureus* is conferred by β-1,4-GlcNAc modified WTA, which requires the glycosyltransferase TarS that is present in nearly all *S. aureus* strains (28,29). Approximately 30% of *S. aureus* strains derived from nasal isolates co-express TarM, a glycosyltransferase that modifies WTA with α-1,4-GlcNAc (28,30). Although α-1,4-GlcNAc did not confer langerin binding, it attenuated langerin binding to β-1,4-GlcNAc WTA, likely as a result of substitution or steric hindrance. This suggests that *S. aureus* clones co-expressing TarM/TarS can alter WTA glycosylation by TarM to evade innate immune activation by LCs (20). Interaction between β-1,4-GlcNAc expressing *S. aureus* and langerin increased pro-inflammatory cytokine production by *in vitro*-generated LCs and in the skin of human langerin-transgenic mice after epicutaneous infection, suggesting a contribution to anti-bacterial host defense (20). Overall, WTA glycosylation impacts the ability of LCs to sense invading *S. aureus* and mount a local immune response.

In addition to TarM and TarS, a third glycosyltransferase, TarP, has recently been identified (31). TarP modifies the WTA backbone with β-linked GlcNAc residues similar to TarS but at the C3 position of RboP instead of C4. *TarP* is always co-expressed with *tarS* and is associated with, but not limited to, healthcare-associated and livestock-associated MRSA strains belonging to clonal complexes 5 and 398 (31,32). TarP can functionally replace TarS with regard to β-lactam resistance and phage susceptibility (31). However, whether the same applies to immune recognition remains to be fully elucidated. For example, TarP-modified WTA displayed attenuated immunogenicity in mice compared to TarS-modified WTA and co-modification of WTA by TarP may lower *S. aureus* antibody recognition despite the presence of antibodies to both WTA glycoforms in serum from healthy individuals (23,32).

In this study, we assessed the impact of TarP-mediated WTA glycosylation on langerin recognition and responses, i.e. antigen uptake and cytokine production, of *in vitro*-generated LCs. We describe that langerin-mediated recognition and uptake of *S. aureus* is similar for strains expressing β-1,3 GlcNAc WTA or β-1,4 GlcNAc WTA. Despite similar recognition and uptake, LC cytokine production was more pronounced upon interaction with *tarP*-expressing bacteria compared to *tarS*-expressing bacteria. Finally, employing synthetic WTA molecules with specific GlcNAc modifications (33), we demonstrate that β-GlcNAc WTA is sufficient but not exclusively required for *S. aureus* binding to langerin-expressing cells. Overall, we provide evidence that LCs are able to sense and respond to all *S. aureus* strains that produce β-GlcNAc-modified WTA. Furthermore, the use of chemically synthesized WTA structures provides a valuable toolbox to study the interaction between host immune molecules such as CLRs and *S. aureus* WTA in more detail.

## Materials & Methods

### Bacterial strains and culture conditions

All plasmids and strains used in this study are listed in Table S1. Bacteria were grown overnight in five ml Todd-Hewitt broth (THB; Oxoid) at 37°C with agitation. Growth medium was supplemented with 10 μg/ml chloramphenicol (Sigma) for plasmid-complemented *S. aureus* strains. Overnight cultures were subcultured the next day in fresh THB and grown to a mid-exponential growth phase, corresponding to an optical density of 0.6-0.7 at 600 nm (OD_600_).

### Generation of complemented N315 Δ*tarS*Δ*tarP* strains

Plasmids containing the shuttle vector RB474 with full-length copies of *tarS* or *tarP* as inserts were isolated from complemented RN4220 Δ*tarM*Δ*tarS* strains (34), and transformed into *Escherichia coli* DC10B by heat shock. Competent *S. aureus* N315 Δ*tarS*Δ*tarP* cells were transformed with pRB474-*tarS* or pRB474-*tarP* (isolated from *E. coli* DC10B) through electroporation with a Bio-Rad Gene Pulser II (100 ohm, 25 μF, 2.5 kV). After recovery, bacteria were plated on Todd-Hewitt agar supplemented with 10 μg/ml chloramphenicol to select plasmid-complemented colonies. The presence of *tarS* or *tarP* was confirmed by PCR analysis, using the primers for TarP (up) 5’-CTTCACGAAAGAGCACTAGAAG-3’ and TarP (dn) 5’-TTCCCGGCAAGTTGGTG-3’ and for TarS (up) 5’-GTGAACATATGAGTAGTGCGTA-3’and TarS (dn) 5’-CATAATGTCCTTCGCCAATCAT-3’. The corresponding WTA glycoform of complemented strains was also verified by bacterial staining with WTA-specific Fab fragments, followed by staining with goat F(ab’)_2_ anti-human kappa-Alexa Fluor 647 (5 μg/ml, Southern Biotech) (Supporting figure 1A).

### Bacterial binding to recombinant human langerin

Bacteria were grown to mid-exponential phase as described above and collected by centrifugation (10 minutes, 4,000 rpm). Supernatant was discarded and bacteria were resuspended to an OD_600_ of 0.4, which corresponds to approximately 10^8^ colony forming units (CFU)/ml in TSM buffer (2.4 g/L Tris (Roche), 8.77 g/L NaCl (Sigma Aldrich), 294 mg/L CaCl_2_.2H20 (Merck), 294 mg/L MgCl_2_.6H20 (Merck), pH=7.4) containing 0.1% bovine serum albumin (BSA, Merck). Next, bacteria were incubated at 37°C for 30 minutes with FITC-labeled human langerin-extracellular domain (ECD) constructs, referred to as human langerin-FITC, as previously described (20,35). Bacteria were washed once with TSM 0.1% BSA, fixed in 1% formaldehyde in PBS and analyzed by flow cytometry on a FACSverse (BD Biosciences). Per sample, 10,000 gated events were collected and data was analyzed using FlowJo 10 (FlowJo, LLC).

### Recombinant expression of monoclonal antibodies and Fab fragments

For monoclonal antibody expression, we cloned the human IgG1 heavy chain (hG) and kappa light chain (hK) constant regions (sequences as present in pFUSE-CHIg-hG1 and pFUSE2-CLIg-hk; Invivogen) in the XbaI-AgeI cloning site of the pcDNA34 vector (Thermo Fisher). VH and VL sequences from monoclonal antibodies specific for α-GlcNAc-WTA (4461), β-GlcNAc-WTA (4497) and β-1,4-GlcNAc-WTA (6292) were derived from patent WO 2014/193722 A1 (36). As the VL of anti-WTA antibody 6292 resulted in precipitation problems, it was adapted towards a Vκ3, leaving the CDR regions (in bold) intact (VL(6292-Vκ3:

EIVLTQSPATLSLSPGERATLSC**RASQGIRNGLG**WYQQKPGQAPRLLIY**PASTLE**SGVPARFSGS GSGTDFTLTISSLEPEDFAVYYC**LQDHNYPP**TFGQGTKVEIK). The VH and VL sequences, preceded by a Kozak sequence (ACCACC) and the HAVT20 signal peptide (MACPGFLWALVISTCLEFSMA) were codon optimized for human expression and ordered as gBlocks (IDT). We cloned VH and VL gBlocks into the pcDNA34 vector, upstream of the IgG1 heavy chain (hG) and kappa light chain (hK) constant regions, respectively, by Gibson assembly (New England Biolabs) according to the manufacturer’s instructions. NheI and BsiWI were used as the 3’ cloning sites for VH and VL, respectively, in order to preserve the immunoglobulin heavy and kappa light chain amino acid sequence. The constructs were transformed in E. coli TOP10F’ by heat shock and clones were verified by PCR and Sanger sequencing (Macrogen). Plasmids were isolated by NucleoBond Xtra Midi kit (Macherey-Nagel) and sterilized using 0.22 μm Spin-X centrifuge columns (Corning). We used EXPI293F cells (Thermo Fisher), grown in EXPI293 Expression medium (Thermo Fisher) at 37°C, 8% CO_2_ in culture filter cap conical flasks (Sigma) on a rotation platform (125 rotations/min) for protein production. One day before transfection, cells were diluted to 2×10^6^ cells/ml and 100 mL cell culture was used for transfection the next day. In 10 mL Opti-MEM (Thermo Fisher), 500 μl PEI-max (1 μg/μl; Polysciences) was mixed with DNA (1 μg/ml cells) in a 3:2 ratio of hK and hG vectors. After 20 minutes incubation at room temperature, this DNA/PEI mixture was added dropwise to 100 ml EXPI293F cells. After five days, we verified IgG expression by SDS-PAGE and harvested cell supernatant by centrifugation and subsequent filtration through a 0.45 μM filter. IgG was purified using a HiTrap Protein A column (GE Healthcare) and Äkta Pure (GE Healthcare). Protein was eluted in 0.1 M citric acid, pH 3.0, and neutralized with 1 M Tris, pH 9.0. The IgG fraction was dialyzed overnight against PBS at 4°C. Purified monoclonal antibodies were stored at −20°C. Fab fragments specific for α-GlcNAc-WTA (4461), β-GlcNAc-WTA (4497) and β-1,4-GlcNAc-WTA (6292) were cloned and expressed similar as the fulllength monoclonal antibodies, except that the Fab heavy chain ends with _211_VEPKSC_216_. A flexible linker (GGGGS), an LPETG and a 6xHIS tag was added at the C-terminus of each Fab. EXPI293F expression supernatant was dialyzed against 50 mM Tris, 500 mM NaCl; pH8.0, before Fab purification on a HISTrap FF column (GE Healthcare). Fab fragments were dialyzed against 50 mM Tris, 300 mM NaCl; pH8.0 and stored at −20°C.

### Production of biotinylated ribitolphosphate (RboP) hexamer (6-) and dodeca (12-)mer

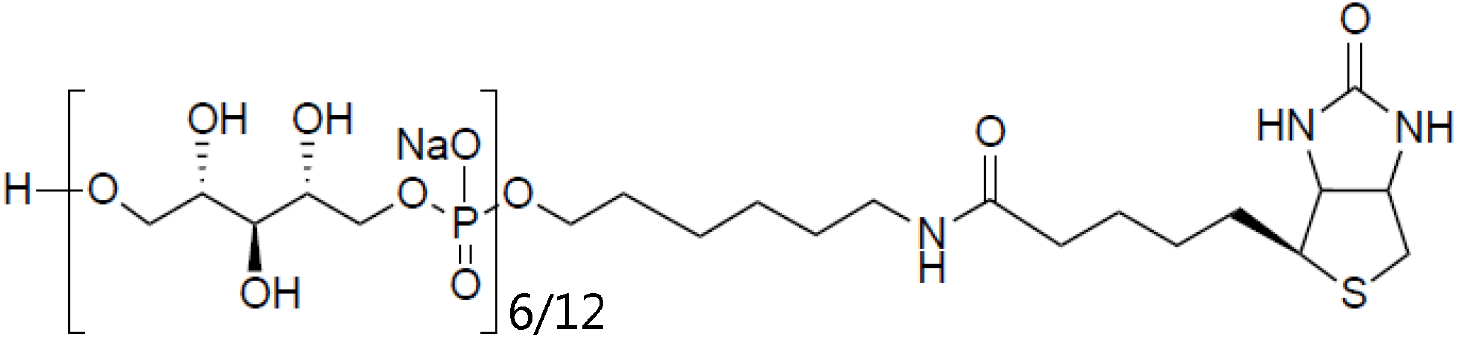

Biotinylated RboP hexamers were synthesized as described previously (23,31). The synthesis of biotinylated RboP dodecamers and chemically defined glycosylated RboP hexamers will be described in detail elsewhere (S. Ali et al, paper in preparation).

### Enzymatic glycosylation of RboP oligomers

Recombinant TarP protein and transformed *E.coli* TOP10F’ strains with pBAD-*tarM* or pBAD-*tarS* were kindly provided by Prof. Thilo Stehle (University of Tübingen, Germany) (31,37). Biotinylated RboP oligomers (0.17 mM) were incubated with recombinant glycosyltransferases TarS, TarP or TarM (6.3 μg/ml) for 2 hours at room temperature with UDP-GlcNAc (2 mM, Merck) in glycosylation buffer (15 mM HEPES, 20 mM NaCl, 1 mM EGTA, 0.02% Tween 20, 10 mM MgCl_2_, 0.1% BSA, pH=7.4). Glycosylated RboP hexamers were coupled to beads by adding 5×10^7^ Dynabeads M280 Streptavidin (Thermo Fisher Scientific) to the individual glycosylation reaction mixtures. After incubation for 15 minutes at room temperature, the coated beads were washed three times with PBS 0.1% BSA 0.05% Tween-20 using a magnetic sample rack and stored at 4°C.

### Recombinant Langerin binding to synthetic WTA

Maxisorb plates (Nunc) were coated with 10 μg/ml his-tetrameric-streptavidin-LPETG overnight at 4°C, which was expressed and isolated from a pColdl-Stav-LPETG vector kindly provided by Tsutomu Tanaka (Kobo University, Japan). The plates were washed three times with TSM 0.05% Tween-20 (TSMT), and subsequently blocked with TSM 1% BSA for 1 hour at 37°C. After three washing steps with TSMT, a 50-fold dilution of the glycosylation mixture described above (corresponding to 0.5 nM RboP 6-mer or 12-mer) was added to the plates and incubated for 1 hour at 37°C. Next, the plates were washed with TSMT, and further incubated with a concentration range of recombinant human langerin-FITC for 30 minutes at 37°C. For blocking experiments, mannan (20 μg/ml) or EGTA (10 mM) were added immediately prior to addition of recombinant human langerin-FITC. Finally, after three washing steps, the plates were analyzed for langerin binding using a Clariostar plate reader (BMG Labtech; excitation 495 nm, emission 535 nm, gain 2,000).

### Cell culture and muLC differentiation

MUTZ-3 cells (ACC-295, DSMZ) were provided by Prof. T. de Gruijl (Amsterdam UMC, The Netherlands). Cells were maintained at a cell density of 0.5 - 1×10^6^ cells/ml in 12-well tissue culture plates (Corning) in MEM-alpha (Gibco) with 20% FBS (Hyclone), 1% glutaMAX (Gibco), 10% spent medium from the renal carcinoma cell line 5637 (ACC-35, DSMZ) and 100 U/ml penicillin-streptomycin (Gibco). Cells were routinely cultured at 37°C with 5% CO_2_. Differentiation of MUTZ-3 cells into MUTZ-3-derived LCs (muLCs) was performed according to described protocols (38,39). In short, MUTZ-3 cells were differentiated in the presence of 100 ng/ml GM-CSF (Genway Biotech), 10 ng/ml TGF-β (R&D systems) and 2.5 ng/ml TNF-α (R&D systems) for 11 days. Twice a week, half of the medium was replaced with fresh medium and double concentration of cytokines. To verify the differentiated muLC phenotype, cells were analyzed by flow cytometry for expression of CD207 (clone DCGM4, Beckman Coulter) and CD1a (clone Hl149, BD Biosciences) as well as the absence of CD34 (clone 581, BD Biosciences).

THP-1 cells, transfected with a lentiviral human langerin construct or empty vector, were cultured in RPMI-1640 (Lonza) supplemented with 10% heat-inactivated FBS and 100 U/ml penicillin-streptomycin (Gibco) as described in (20).

### Binding and internalization of WTA beads or *S. aureus* by Langerin-expressing cells

Dynabeads-M280 Streptavidin (Thermo Fisher Scientific) and mid-exponential *S. aureus* (OD_600_= 0.6-0.7) were labeled with 0.5 mg/ml FITC (Sigma) in PBS for 30 minutes at 4°C. After extensive washing and coating of the beads with glycosylated RboP hexamers as described above, beads and bacteria were resuspended in RPMI 0.1% BSA at a concentration of 5×10^7^ beads/ml or 1×10^8^ CFU/ml (OD_600_= 0.4), respectively. Bacteria were stored at −20°C and beads at 4°C in the dark. For binding experiments, 1×10^5^ cells (THP-1 cells or muLCs) were incubated with FITC-labeled WTA beads or FITC-labeled *S. aureus* at different ratios in RPMI 0.1% BSA for 30 minutes at 4°C. Cells were washed (300 x *g* for 10 minutes at 4°C), fixed in PBS 1% formaldehyde and analyzed by flow cytometry as described above. To quantify internalization of β-GlcNAc WTA beads by THP-1 cells, we incubated WTA beads with 2×10^5^ cells in RPMI 0.1% BSA at a bead-to-cell ratio of 1 for 30 minutes at 4°C. Cells were washed twice to remove unbound beads, and the sample was divided over two separate tubes. Both samples were incubated for an additional 30 minutes, one at 4°C and the other at 37°C with 5% CO_2_ to allow phagocytosis. Cells were washed, and Fc-receptors were blocked with recombinant FLIPR-like (6 μg/ml) for 15 minutes at 4°C (40). Next, monoclonal antibodies specific for β-GlcNAc or α-GlcNAc WTA (4497/4461-IgG1, respectively) were added to all samples at 3 μg/ml for 20 minutes at 4°C, followed by goat anti-human kappa-Alexa Fluor 647 (5 μg/ml, Southern biotech) for another 20 minutes at 4°C to allow discrimination between cell adherent (FITC+/Alexa fluor 647+) and internalized beads (FITC+/Alexa fluor 647-). Finally, cells were washed and fixed in 1% formaldehyde in PBS. The internalized fraction was calculated from the loss of Alexa Fluor 647 signal of FITC+ cells by flow cytometry, as previously described (41).

To confirm bead internalization by confocal microscopy, cells were stained with WGA-Alexa Fluor 647 (Thermo Fisher Scientific) and DAPI (Sigma) following incubation for 30 minutes at 37°C with FITC-labeled WTA beads and coated on 8 well chamber slides glass slides (Ibidi) before analysis by confocal laser scanning microscopy (SP5, Leica).

### muLC stimulation

Gamma-irradiation of *S. aureus* and stimulation of muLCs was performed as previously described (20). Briefly, *S. aureus* strains were grown to exponential phase, washed with PBS, concentrated 10-fold in PBS with 17% glycerol, and stored at −80°C. Gamma-irradiation of bacteria was performed at Synergy Health Ede B.V., a STERIS company (Ede, The Netherlands). The loss of viability was confirmed by plating, and the bacterial concentrations were calculated using the MACSQuant Analyzer 10.

1×10^5^ muLCs were stimulated with γ-irradiated RN4220 Δ*tarMS*+*ptarS*, RN4220 Δ*tarMS*+*ptarP* or RN4220 Δ*tarMS*+*ptarM* at bacteria to cell ratios of 0, 1, 10, and 50 for 24 hours at 37°C with 5% CO_2_ in IMDM containing 10% FBS. Supernatants for cytokine analysis were collected after centrifugation (300 x g, 10 min at 4°C), and stored at −80°C until further analysis. Cells were washed with PBS 0,1% BSA, stained with CD83 (clone HB15e) and CD86 (clone IT2.2, Sony Biotechnology), fixed and analyzed by flow cytometry. Cytokine production was analyzed by ELISA, for IL-8 (Sanquin) and TNFα (Thermo Fisher) following manufacturer’s instructions.

### Statistical analysis

Flow cytometry data was analyzed using FlowJo 10 (FlowJo, LLC). All data was analyzed using GraphPad Prism 8.3 (GraphPad Software), with a Two-way ANOVA followed by a Dunnett’s multiple comparison test except for bacterial binding to langerin-FITC at one fixed concentration for which oneway ANOVA was performed with Dunnett’s multiple comparison test. P values are depicted in the figures and *p* < 0.05 was considered significant.

## Results

### TarP and TarS both confer binding of human langerin to *S. aureus*

TarP can replace several key functions of TarS, including resistance to β-lactam antibiotics and susceptibility to siphophage infection (31). In contrast, decoration of WTA with β-1,3-GlcNAc in addition to or instead of β-1,4-GlcNAc may impact immune detection by antibodies (23,31). We recently identified that β-1,4-GlcNAc WTA is specifically detected by the human innate receptor langerin (20). To assess whether human langerin was also able to detect *tarP*-expressing *S. aureus* strains, we employed a FITC-labeled recombinant construct of the extracellular carbohydrate domain (ECD) of human langerin (langerin-FITC) (35). Using *S. aureus* strain N315 that naturally expresses both *tarS* and *tarp* (31), we observed that langerin binding was only significantly impaired upon deletion of both glycosyltransferases (Δ*tarPS*), but not in either of the single mutant strains (Fig. 1A). Subsequent complementation of the Δ*tarPS* double mutant with a plasmid containing either *tarS* or *tarP* restored the binding to recombinant langerin-FITC (Fig. 1A). This observation in differential langerin binding amongst the N315 mutant panel persisted over a 100-fold concentration range of langerin-FITC, although at higher concentrations langerin-FITC also showed significant binding to the Δ*tarPS* strain (Fig. 1B). Binding to the N315 Δ*tarPS* strain was also dependent on the langerin CRD since the interaction could be blocked by addition of mannan (Supporting Fig. 1B). Similar binding experiments were additionally performed in *S. aureus* strain RN4220, which naturally co-expresses *tarS* and *tarM*, but not *tarP*. As previously reported (20), langerin binding to RN4220 wild-type was significantly reduced in the Δ*tarMS* double mutant (Fig. 1C). Binding could be restored by complemention with either *tarS* or *tarP* but not *tarM* (Fig. 1C). For the N315 and RN4220 strain panels, expression of the correct WTA glycoform was confirmed through binding of specific mAbs (Supporting Fig. 1A;(23)). Overall, langerin binds to TarP-modified WTA independent of strain background.

**Figure 1.**
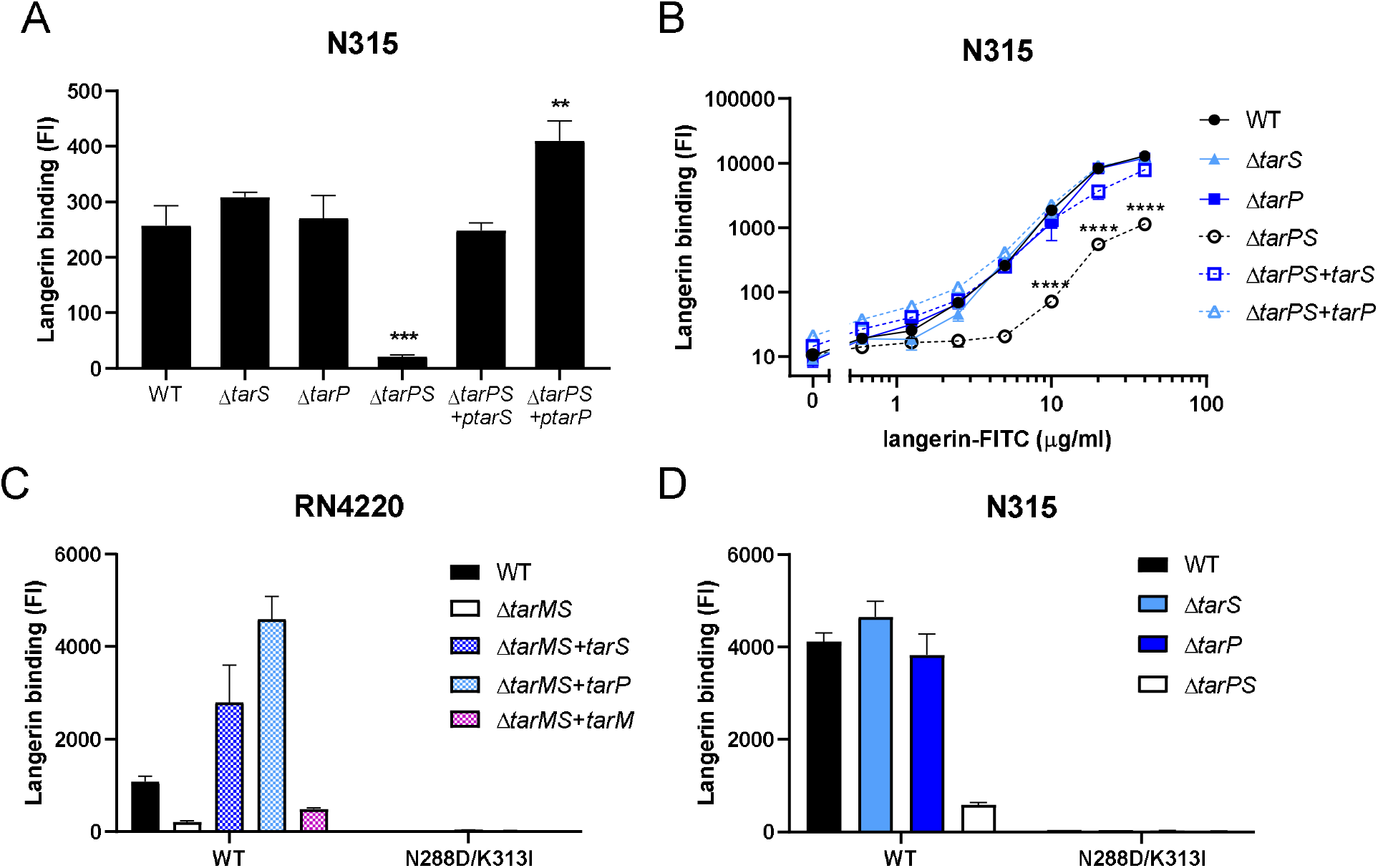
WTA β-GlcNAcylation by TarS and TarP confers langerin binding to *S. aureus*. Binding of recombinant human langerin-FITC A) to N315 WT, Δ*tarS*, Δ*tarP*, Δ*tarPS*, Δ*tarPS* +*ptarS* and Δ*tarPS* +*ptarP* at a fixed concentration of 5 μg/ml, B) to the indicated N315 strain panel using a concentration range of langerin-FITC (0.6-40 μg/ml). C, D) Binding of FITC-labeled recombinant human langerin wildtype and N288D/K313I double SNP variant (10 μg/ml) to C) RN4220 WT, Δ*tarMS*, Δ*tarMS* +*ptarS*, Δ*tarMS* +*ptarP* and Δ*tarMS* +*ptarM* and D) the N315 mutant panel (mentioned above). Data is depicted as geometric mean fluorescence intensity (FI) + standard error of mean (SEM) of biological triplicates. \*\**p* < 0.01, \*\*\**p* < 0.001, \*\*\*\**p* < 0.0001.

While it was apparent that langerin binding to *S. aureus* required either TarP or TarS, it was not clear whether the receptor bound the two different epitopes in a similar way. Previously, we showed that langerin binding to *S. aureus* was abrogated when a naturally-occurring double SNP was introduced into the human langerin ECD (41). Using these same langerin SNP constructs, we observed a similar loss of binding to TarP-expressing *S. aureus* (Fig. 1C, D). These data suggest that the WTA β-1,3-GlcNAc epitope created by TarP is similarly dependent on these two residues in the carbohydrate recognition domain of langerin compared to the β-1,4-GlcNAc epitope on WTA generated by TarS.

### WTA β-GlcNAc is sufficient to confer langerin binding

TarP-expressing *S. aureus* can bind langerin in a similar way to *S. aureus* expressing TarS. However, we also observed significant residual binding in the Δ*tarPS* background at higher langerin concentrations (Fig. 1B). We therefore asked whether WTA-β-GlcNAc is sufficient to confer binding to *S. aureus* or whether additional bacterial co-factors are required. The isolation of WTA from the bacterial cell wall is challenging; the procedure is labor intensive but moreover, the instability and variation in isolated WTA creates difficulties for assay reproducibility. Therefore, we used our previously developed system (23), where chemically-synthesized WTA backbone fragments of defined length are glycosylated by specific recombinant Tar enzymes *in vitro* (Fig. 2A). With this robust system, we have previously studied the interaction of specific WTA glycoforms and antibodies in a reproducible and low background manner(23). In this study, we used both hexameric and dodecameric RboP backbones to assess the influence of WTA chain length on langerin binding. Differently glycosylated biotinylated WTA structures were coated on streptavidin-coated ELISA plates and incubated with a concentration range of recombinant langerin-FITC. Only wells coated with β-1,4-GlcNAc- and β-1,3-GlcNAc-glycosylated WTA structures mediated concentration-dependent binding to langerin and no binding was observed to the RboP backbone or α-1,4-GlcNAc-glycosylated WTA (Fig. 2B, C). In addition, langerin binding was increased when the WTA backbone was extended from 6- to 12-RboP units (Fig. 2B, C). Interaction between recombinant langerin-FITC and synthetic WTA was completely abolished in the presence of EGTA (Fig. 2C), which scavenges calcium ions required for receptor binding. Langerin binding likely requires more than two β-GlcNAc residues, since we could not detect binding to a fully synthetic WTA molecule consisting of hexameric RboP backbone and β-1,4-GlcNAc coupled to the third and terminal RboP subunit (Supporting Fig 2A, B). In contrast, monoclonal antibodies specific for either α-GlcNAc-WTA or β-GlcNAc-WTA were able to bind the fully synthetic WTA structures (Supporting Fig 2C). This does not only indicate that fully-synthetic structures were coated correctly to the wells but also underlines the differences in minimal binding requirements to glycosylated WTA between antibodies and langerin. Overall, these data confirm that β-GlcNAc WTA is sufficient to confer interaction with langerin and does not require the presence of D-alanine residues on WTA nor additional bacterial factors.

**Figure 2.**
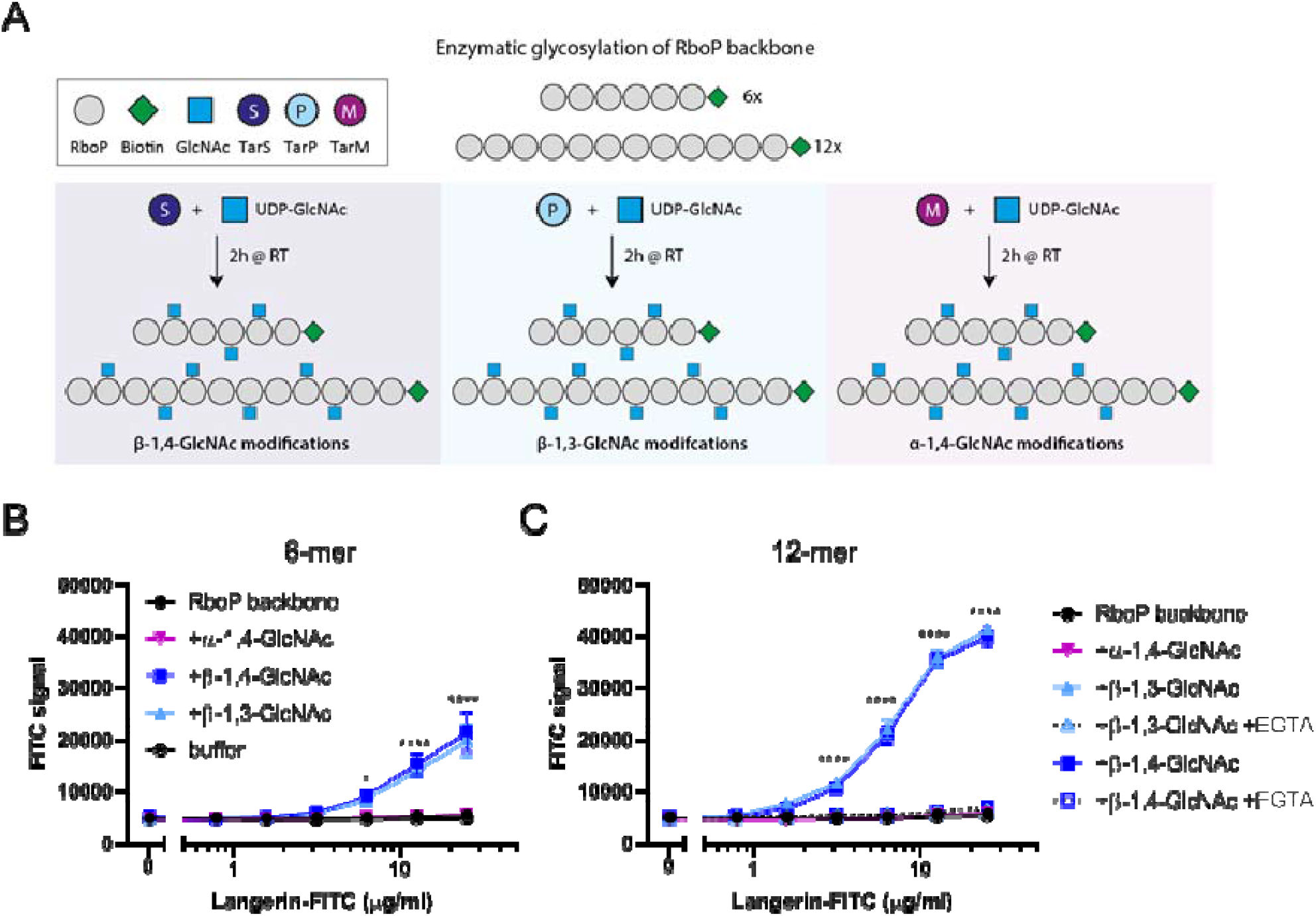
β-GlcNAc-modified WTA is sufficient to confer langerin binding. A) Schematic overview of the synthetic WTA structures and *in vitro* glycosylation by recombinant TarS, TarP or TarM. B) Binding of recombinant human langerin-FITC (0.4-25 μg/ml) to RboP hexamers alone (RboP backbone) or after *in vitro* glycosylation by TarS, TarP or TarM. C) Binding of recombinant human langerin-FITC (0.4-25 μg/ml) to RboP dodecamers alone (RboP backbone) or after *in vitro* glycosylation similar to RboP hexamers. Binding was assessed in the absence and presence of EGTA (10 mM). Data for panel B and C is shown as fluorescence signal + SEM of three independent experiments. \**p* < 0.05, \*\*\*\**p* < 0.0001.

We also assessed binding of beads, coated with the differently glycosylated WTA oligomers, to surface-expressed langerin on transfected THP-1 cells. FITC-labeled beads were coated with synthetic glycosylated WTA hexamers, and coating was verified by binding of monoclonal antibodies specific for either α-GlcNAc or β-GlcNAc WTA (Supporting Fig. 3). We observed strong binding of β-GlcNAc WTA beads, modified by either TarS or TarP, to langerin-expressing THP-1 cells but not empty vector control cells (Fig. 3A). In addition to binding, Langerin+ THP-1 cells internalized the majority of adhered beads as assessed by flow cytometry (Fig. 3B) and confocal microscopy (Fig. 3C). No apparent differences in receptor binding or cellular uptake were observed for TarS- and TarP-modified WTA beads in this system, suggesting that both epitopes confer a similar function with regard to langerin interaction.

**Figure 3.**
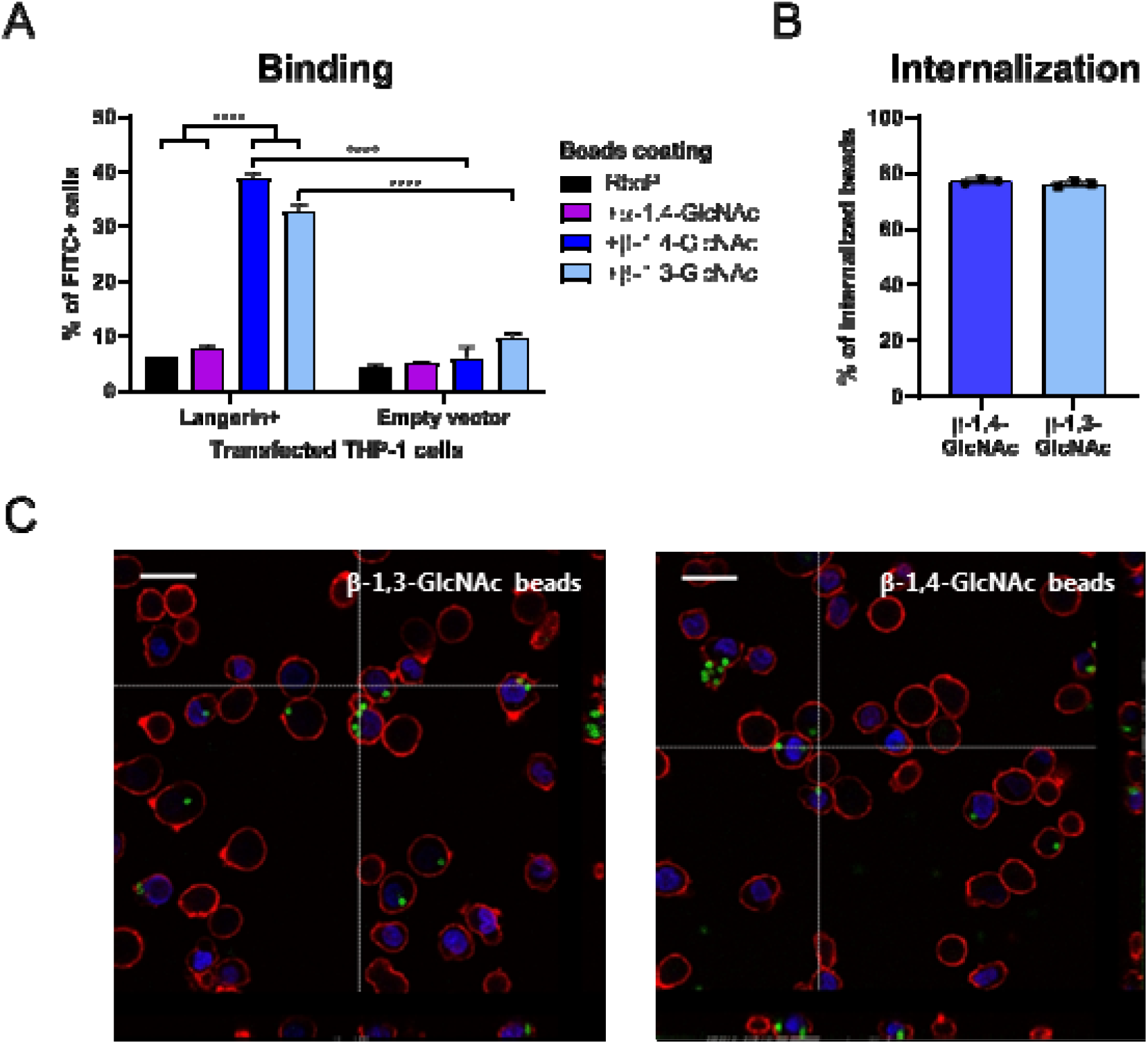
Binding and internalization of β-GlcNAc-WTA-coated beads by langerin-expressing THP-1 cells. A) Binding of FITC-labeled beads, coated with unglycosylated or *in vitro* glycosylated RboP hexamers, to THP-1 cells transfected with human langerin or empty vector at a bead-to-cell ratio of 1. Adherence is represented by % of FITC+ cells. B) Proportion of adherent β-GlcNAc WTA beads that is internalized by Langerin+ THP-1 cells. C) Confocal microscopy images (40X) of β-GlcNAc WTA beads (FITC-labeled: green) bound to and internalized by Langerin+THP-1 cells (WGA-Alexa 647: red, DAPI: blue). Vertical lines correspond to cross section of z-stack on the right, horizontal lines to cross section below, scale bars correspond to 25 μm. For panels A and B, graphs represent mean + SEM of biological triplicates, \*\*\*\**p* < 0.000

### The expression of β-GlcNAc WTA contributes significantly to the interaction between *S. aureus* and LCs

We have recently shown that langerin significantly contributes to the interaction between *S. aureus* and primary human LCs (20). In addition, *in vitro*-generated muLCs were used as a LC cell model to demonstrate the impact of langerin recognition on activation of APCs (20). Here, we again used muLCs to study the binding of surface-expressed langerin to β-GlcNAc WTA modifications mediated by TarS or TarP. In line with the THP-1 binding experiments, muLCs also specifically bound to β-GlcNAc WTA beads, irrespective of linkage to C3 (TarP) or C4 (TarS) (Fig. 4A). At a bead-to-cell ratio of 10, beads decorated with β-1,3-GlcNAc WTA adhered significantly better compared to beads decorated with β-1,4-GlcNAc WTA (Fig. 4A). This observed binding was mediated by the presence of langerin, as we were able to block the binding of muLCs to β-GlcNAc WTA beads by addition of mannan, a ligand for langerin, or specific langerin-blocking monoclonal antibodies (Fig. 4B). These data show that β-GlcNAcylated WTA is sufficient to confer binding to muLCs and does not require bacterial co-factors.

**Figure 4.**
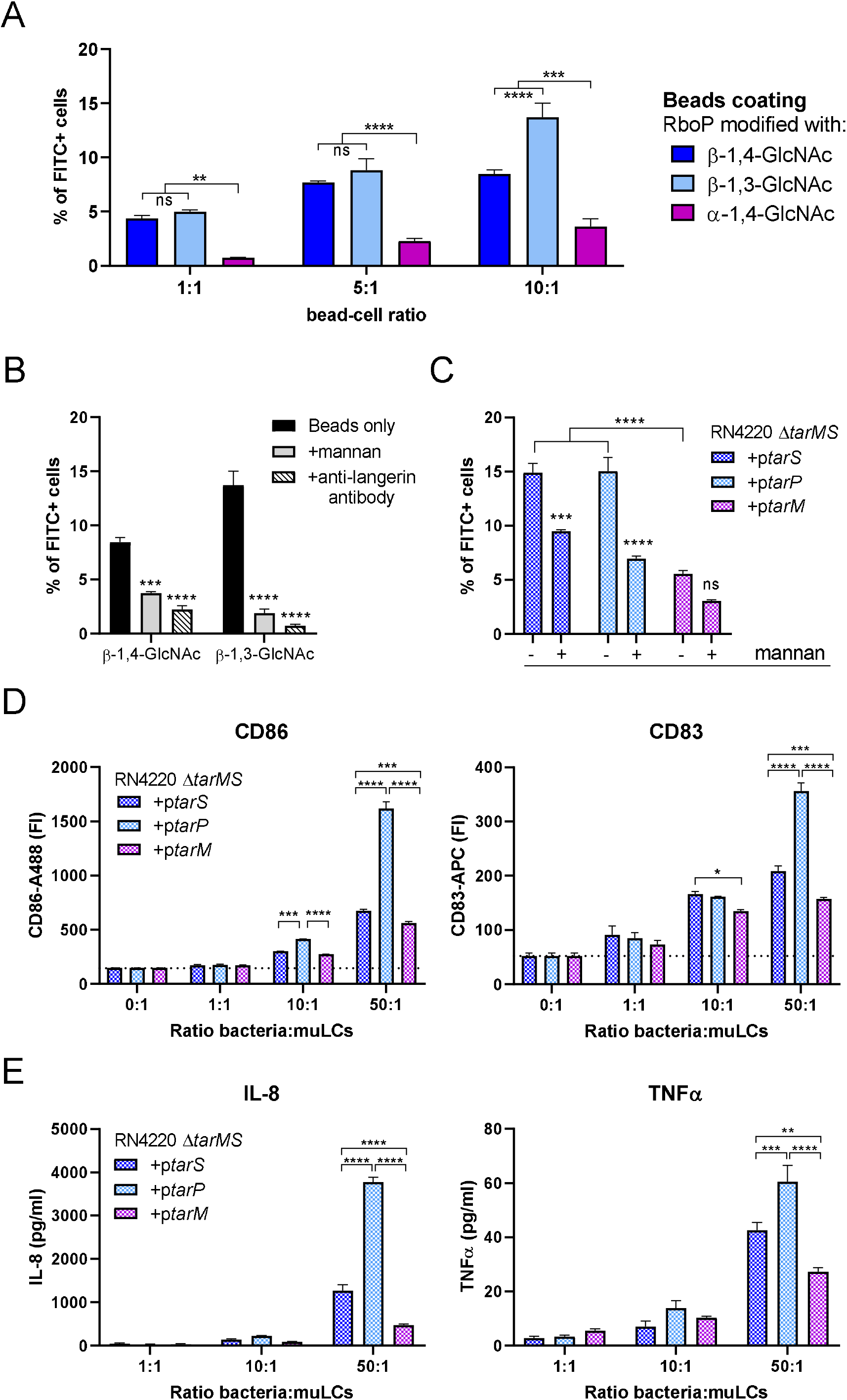
*S. aureus* WTA glycoform affects binding to and activation of *in vitro*-generated LCs. A) Binding of FITC-labeled beads, coated with *in vitro* glycosylated RboP dodecamers, to muLCs at bead-to-cell ratios of 1, 5, and 10. Bead adherence is displayed as % of FITC+ cells. B) Binding of FITC-labeled beads coated with TarS- or TarP-modified RboP dodecamers to muLCs at a bead-to-cell ratio of 10 in the absence (similar to A) or presence of mannan (20 μg/ml) or anti-langerin blocking antibody (20 μg/ml). C) Binding of FITC-labeled RN4220 Δ*tarMS* complemented with plasmid-expressed *tarS, tarP* or *tarM* to muLCs at a bacteria-to-cell ratio of 1. Bacterial binding is represented by % of FITC+ cells D) Surface expression of activation marker CD86 and maturation marker CD83 by muLCs after 24h stimulation with γ-irradiated RN4220 Δ*tarMS* complemented with plasmid-expressed *tarS, tarP* or *tarM*,at bacteria-to-cell ratios of 1, 10, and 50. E) Concentration of IL-8 and TNFα in the supernatant of muLCs described in D. The data for all panels represent mean + SEM of biological triplicates. \**p* < 0.05, \*\**p* < 0.01, \*\*\**p* < 0.001, \*\*\*\**p* < 0.0001.

Next, we assessed whether β-GlcNAc WTA was necessary for *S. aureus* binding to muLCs. For these experiments we used the RN4220 Δ*tarMS* background where *tarM, tarS* and *tarP* are individually and constitutively expressed from a complementation plasmid. We observed an approximately 3-fold higher binding to muLCs by *S. aureus* strains expressing β-GlcNAc WTA compared to α-GlcNAc-WTA producing *S. aureus* (Fig. 4C). However, even in the absence of β-GlcNAc WTA, *S. aureus* was able to adhere to muLCs. Furthermore, binding of β-GlcNAc WTA producing *S. aureus*, but not α-GlcNAc producing *S. aureus*, to muLCs was significantly blocked by addition of mannan (Fig. 4C). These results indicate that the interaction between Langerin and β-GlcNAc WTA is an important determinant, although not exclusively required, for *S. aureus* binding to LCs.

To assess the downstream effects of langerin-mediated binding of *S. aureus* to muLCs, and potential differences herein between β-1,4-GlcNAc-WTA versus β-1,3-GlcNAc-WTA producing *S. aureus*, we stimulated muLCs for 24 hours with gamma-irradiated RN4220 Δ*tarMS*, complemented with either plasmid-expressed *tarS, tarP* or *tarM*. Surface expression of activation markers CD86 and CD83 increased in a dose-dependent manner in response to all three strains. Expression of CD86 and CD83 was highest in response to *tarP*-complemented *S. aureus* and differed significantly from *tarS*-complemented *S. aureus* (Fig. 4D). The production of IL-8 and TNF-α showed a similar pattern, where all three strains induced a dose-dependent cytokine response with highest cytokine levels in response to *tarP*-complemented *S. aureus* (Fig. 4E). In line with previous results, *tarM*-complemented *S. aureus* showed the lowest activation of muLCs, both in surface expression of CD86 and CD83, as well as cytokine production. This data suggests that besides the known effect between α-GlcNAc-WTA and β-GlcNAc-WTA, there could be additional differences in langerin-mediated LC activation between β-1,3-GlcNAc-WTA and β-1,4-GlcNAc-WTA.

## Discussion

This study expands our previous findings that human LCs sense the major skin pathogen *S. aureus*.Thereby, LCs may contribute to host defense when *S. aureus* invades the skin through early initiation of pro-inflammatory responses and recruitment of neutrophils. We show here that the CLR langerin can detect *S. aureus* strains that produce β-GlcNAcylated WTA, which can be mediated by the housekeeping glycosyltransferase TarS and the accessory enzyme TarP (20,31). Using a combination of recombinant langerin and langerin-transfected cell lines, genetically-modified *S. aureus* strains and *in vitro* generated LCs, we suggest that the interaction between langerin and *tarP*-expressing *S. aureus* results in similar binding but quantitatively-different immunological responses. Moreover, comparing the binding of beads coated with synthetic glycosylated WTA oligomers and *S. aureus* modified strains emphasizes that the interaction between LCs and *S. aureus* is largely, but not solely, dependent on the expression of β-GlcNAc WTA.

Binding of recombinant langerin to *S. aureus* was abrogated in bacteria that lack WTA glycosyltransferases, i.e. *N315* Δ*tarPS* and RN4220Δ*tarMS* bacteria. However, at higher concentrations, residual langerin binding to these WTA-deglycosylated strains was still observed, suggesting the presence of a second, currently unidentified minor ligand for langerin on the *S. aureus* surface. This observed binding was specific, as the binding was saturable and was inhibited by addition of mannan (Supporting Fig. 1B). *S. aureus* expresses a wide variety of surface proteins that contribute to skin colonization and infection (42). Interestingly, some of these proteins, such as the serine-aspartate repeat (SDR) proteins and SraP, are heavily glycosylated (43-45), thereby representing potential targets for langerin in addition to β-GlcNAc WTA.

The toolbox of synthetic WTA fragments allowed us to gain more insights into the binding requirements of langerin to WTA. Following current consensus, the WTA backbone consists of up to 40 repeating units of RboP that can be co-decorated with D-alanine and GlcNAc residues (46). The synthetic RboP polymers used here are only modified with GlcNAc and do not contain D-alanine residues. Consequently, we conclude that D-alanylation of WTA is dispensable for langerin binding in our assays, although we cannot rule out that the interaction would be affected by the presence of D-alanine. Also, when expressed by *S. aureus*, the absence or presence of D-alanine does not seem to impact langerin binding (Supplementary Fig. 1C). In addition, we observed a strong impact of epitope abundance on langerin binding; doubling the length of the synthetic WTA backbone enhanced langerin binding, which is most likely explained by an increased number of GlcNAc epitopes following *in vitro* glycosylation. Furthermore, we did not observe langerin binding to fully defined WTA structures, which only contained two β-GlcNAc modifications (Supporting Fig. 2B). This could be due to a limited sensitivity of our assay. Alternatively, it may indicate that langerin requires more than two β-GlcNAc epitopes or differently spaced β-GlcNAc moieties to interact. In contrast, two GlcNAc moieties are sufficient for antibodies to interact with WTA (Supporting Fig. 2C). Currently, not much is known about the regulation of WTA biosynthesis and glycosylation, although both the length of the WTA backbone as well as the expression of glycosyltransferases are believed to be affected by environmental cues. In the skin, activation of the Agr regulon results in increased WTA expression on the surface (47). Additionally, TarS-mediated WTA glycosylation increases under infection conditions at the expense of TarM- or TarP-mediated glycosylation, which dominate WTA glycosylation under *in vitro* growth conditions (31,48,49). Consequently, more β-1,4-GlcNAc epitopes are produced *in vivo* (48), which would greatly enhance receptor avidity of langerin and impact its function (14).

TarP can replace TarS in several key processes, including β-lactam resistance (31). However, whether the same applies to immune recognition still remains to be fully clarified. In mice, TarP-modified WTA appeared less immunogenic as compared to TarS-modified WTA (31). Previous work has shown the existence of cross-reactive human antibodies to both β-GlcNAc epitopes, while other antibodies seem to be more exclusively directed towards β-1,4-GlcNAc (23). Until now, no studies have assessed the potential discrimination between TarS- and TarP-expressing *S. aureus* strains by innate immune cells. From our cell-based assays, β-1,3-GlcNAc-modified WTA has a similar ability to bind langerin compared to β-1,4-GlcNAc-modified WTA. However, LC activation as detected by cytokine production appears to be higher in response to TarP-versus TarS-expressing *S. aureus* strains. This finding potentially underlines an important difference in the stimulatory capacity of both modifications, where β-1,3-GlcNAc is more immunostimulatory for innate responses, whereas β-1,4-GlcNAc is dominant for adaptive antibody recognition. One explanation for this could be the difference in glycosylation between both glycosyltransferases. TarP modifies the RboP backbone with GlcNAc moieties at a higher efficiency than TarS, which could subsequently enhance receptor clustering and internalization by LCs. Moreover, glycosylation by TarS or TarP differentially affects D-alanylation of WTA, resulting in overall charge differences (31). As a consequence, TarP-mediated glycosylation might negatively affect antigen-presentation by APCs as a result of decreased zwitterionic charge properties. As a result, T cell responses and T cell-dependent B cell responses to TarP-modified WTA may be hampered. Furthermore, T cell-independent B cell responses to TarP-modified WTA could be affected as well, via decreased crosslinking of the B cell receptor. However, more research is needed to support this hypothesis, and the synthesis of WTA oligomers with added D-alanine modifications will serve as an excellent tool to study this.

Our results underline the ability of muLCs to detect and internalize *S. aureus* that express β-GlcNAc on their surface. In line with previous work, we observed that langerin recognition increased surface expression of activation markers CD86 and CD83 and enhanced the production of pro-inflammatory cytokines such as IL-8, which would generally serve to recruit neutrophils to the site of infection to promote rapid eradication of invading *S. aureus* (20). Whether and how the interaction between langerin and WTA would contribute to LC-mediated immunity against *S. aureus* still remains to be elucidated. Besides processes such as antigen uptake and presentation to CD4 T cells, little is known about direct downstream responses of langerin (17-19,50). Moreover, a lack of robust models, including limited access to human skin explants, differences in langerin ligand specificity (16) and immune cell subsets in commonly used experimental animals (51), represent significant challenges to study immature LC function. The synthetic WTA oligomers used here could represent a robust tool to specifically study downstream effects of langerin receptor binding, and could even be used in combination with appropriate TLR stimulation to unravel LC responses in response to specific langerin-TLR triggers (52).

Overall, langerin senses all β-GlcNAc WTA-producing *S. aureus* strains, which contributes to but is not exclusively required for recognition by LCs. In addition, we suspect the existence of a second langerin ligand on the surface of *S. aureus*. It is currently difficult to dissect the functional consequences of LCs responses in more relevant biological systems. In addition, we also lack knowledge on *in vivo* expression of WTA glycosyltransferases, the resulting WTA glycoform and the spatial distribution across the bacterial cell wall, which all impact interaction and responses triggered by CLRs such as langerin. Future research will need to elucidate the impact of the *S. aureus* WTA glycoform on the ability of LCs *in situ* to sense invading *S. aureus* in the skin, a frequent point of entry, and whether this interaction aids in prevention of bacterial dissemination by mounting an effective local response.

## Data availability

Upon request, the data supporting these findings are available from the corresponding author.

## Acknowledgements

We thank Dani Heesterbeek and Lisanne de Vor for technical assistance with confocal microscopy.

## Funding and additional information

This work was supported by the European Union’s Horizon 2020 research and innovation program under the Marie Skłodowska-Curie grant agreement No 675106 coordinated by Dr. Fabio Bagnoli (GSK, Siena, Italy) and by a Vidi grant (91713303) from the Dutch Scientific Organization (NWO) to N.M.v.S.

## References

1. Ray, G. T., Suaya, J. A., and Baxter, R. (2013) Microbiology of skin and soft tissue infections in the age of community-acquired methicillin-resistant *Staphylococcus aureus*. Diagn Microbiol Infect Dis 76, 24–30

2. Bagnoli, F., Bertholet, S., and Grandi, G. (2012) Inferring reasons for the failure of *Staphylococcus aureus* vaccines in clinical trials. Front Cell Infect Microbiol 2, 16

3. Doebel, T., Voisin, B., and Nagao, K. (2017) Langerhans Cells - The Macrophage in Dendritic Cell Clothing. Trends in immunology 38, 817–828

4. Seneschal, J., Clark, R. A., Gehad, A., Baecher-Allan, C. M., and Kupper, T. S. (2012) Human epidermal Langerhans cells maintain immune homeostasis in skin by activating skin resident regulatory T cells. Immunity 36, 873–884

5. Sparber, F., De Gregorio, C., Steckholzer, S., Ferreira, F. M., Dolowschiak, T., Ruchti, F., Kirchner, F. R., Mertens, S., Prinz, I., Joller, N., Buch, T., Glatz, M., Sallusto, F., and LeibundGut-Landmann, S. (2019) The Skin Commensal Yeast Malassezia Triggers a Type 17 Response that Coordinates Anti-fungal Immunity and Exacerbates Skin Inflammation. Cell Host Microbe 25, 389–403

6. de Witte, L., Nabatov, A., Pion, M., Fluitsma, D., de Jong, M. A., de Gruijl, T., Piguet, V., van Kooyk, Y., and Geijtenbeek, T. B. (2007) Langerin is a natural barrier to HIV-1 transmission by Langerhans cells. Nat Med 13, 367–371

7. Igyarto, B. Z., Haley, K., Ortner, D., Bobr, A., Gerami-Nejad, M., Edelson, B. T., Zurawski, S. M., Malissen, B., Zurawski, G., Berman, J., and Kaplan, D. H. (2011) Skin-resident murine dendritic cell subsets promote distinct and opposing antigen-specific T helper cell responses. Immunity 35, 260–272

8. Kobayashi, T., Glatz, M., Horiuchi, K., Kawasaki, H., Akiyama, H., Kaplan, D. H., Kong, H. H., Amagai, M., and Nagao, K. (2015) Dysbiosis and *Staphylococcus aureus* Colonization Drives Inflammation in Atopic Dermatitis. Immunity 42, 756–766

9. de Jong, M. A., Vriend, L. E., Theelen, B., Taylor, M. E., Fluitsma, D., Boekhout, T., and Geijtenbeek, T. B. (2010) C-type lectin Langerin is a beta-glucan receptor on human Langerhans cells that recognizes opportunistic and pathogenic fungi. Mol Immunol 47, 1216–1225

10. van der Aar, A. M., Sylva-Steenland, R. M., Bos, J. D., Kapsenberg, M. L., de Jong, E. C., and Teunissen, M. B. (2007) Loss of TLR2, TLR4, and TLR5 on Langerhans cells abolishes bacterial recognition. J Immunol 178, 1986–1990

11. Flacher, V., Bouschbacher, M., Verronese, E., Massacrier, C., Sisirak, V., Berthier-Vergnes, O., de Saint-Vis, B., Caux, C., Dezutter-Dambuyant, C., Lebecque, S., and Valladeau, J. (2006) Human Langerhans cells express a specific TLR profile and differentially respond to viruses and Gram-positive bacteria. J Immunol 177, 7959–7967

12. Brown, G. D., Willment, J. A., and Whitehead, L. (2018) C-type lectins in immunity and homeostasis. Nat Rev Immunol 18, 374–389

13. Valladeau, J., Ravel, O., Dezutter-Dambuyant, C., Moore, K., Kleijmeer, M., Liu, Y., Duvert-Frances, V., Vincent, C., Schmitt, D., Davoust, J., Caux, C., Lebecque, S., and Saeland, S. (2000) Langerin, a novel C-type lectin specific to Langerhans cells, is an endocytic receptor that induces the formation of Birbeck granules. Immunity 12, 71–81

14. Feinberg, H., Powlesland, A. S., Taylor, M. E., and Weis, W. I. (2010) Trimeric structure of langerin. J Biol Chem 285, 13285–13293

15. Feinberg, H., Taylor, M. E., Razi, N., McBride, R., Knirel, Y. A., Graham, S. A., Drickamer, K., and Weis, W. I. (2011) Structural basis for langerin recognition of diverse pathogen and mammalian glycans through a single binding site. Journal of molecular biology 405, 1027–1039

16. Hanske, J., Schulze, J., Aretz, J., McBride, R., Loll, B., Schmidt, H., Knirel, Y., Rabsch, W., Wahl, M. C., Paulson, J. C., and Rademacher, C. (2017) Bacterial Polysaccharide Specificity of the Pattern Recognition Receptor Langerin Is Highly Species-dependent. J Biol Chem 292, 862–871

17. Thepaut, M., Valladeau, J., Nurisso, A., Kahn, R., Arnou, B., Vives, C., Saeland, S., Ebel, C., Monnier, C., Dezutter-Dambuyant, C., Imberty, A., and Fieschi, F. (2009) Structural studies of langerin and Birbeck granule: a macromolecular organization model. Biochemistry 48, 2684–2698

18. Mc Dermott, R., Ziylan, U., Spehner, D., Bausinger, H., Lipsker, D., Mommaas, M., Cazenave, J. P., Raposo, G., Goud, B., de la Salle, H., Salamero, J., and Hanau, D. (2002) Birbeck granules are subdomains of endosomal recycling compartment in human epidermal Langerhans cells, which form where Langerin accumulates. Mol Biol Cell 13, 317–335

19. van der Vlist, M., de Witte, L., de Vries, R. D., Litjens, M., de Jong, M. A., Fluitsma, D., de Swart, R. L., and Geijtenbeek, T. B. (2011) Human Langerhans cells capture measles virus through Langerin and present viral antigens to CD4(+) T cells but are incapable of crosspresentation. Eur J Immunol 41, 2619–2631

20. van Dalen, R., De La Cruz Diaz, J. S., Rumpret, M., Fuchsberger, F. F., van Teijlingen, N. H., Hanske, J., Rademacher, C., Geijtenbeek, T. B. H., van Strijp, J. A. G., Weidenmaier, C., Peschel, A., Kaplan, D. H., and van Sorge, N. M. (2019) Langerhans Cells Sense *Staphylococcus aureus* Wall Teichoic Acid through Langerin To Induce Inflammatory Responses. mBio 10, 1–14

21. Winstel, V., Xia, G., and Peschel, A. (2014) Pathways and roles of wall teichoic acid glycosylation in *Staphylococcus aureus*. Int J Med Microbiol 304, 215–221

22. Lehar, S. M., Pillow, T., Xu, M., Staben, L., Kajihara, K. K., Vandlen, R., DePalatis, L., Raab, H., Hazenbos, W. L., Hiroshi Morisaki, J., Kim, J., Park, S., Darwish, M., Lee, B. C., Hernandez, H., Loyet, K. M., Lupardus, P., Fong, R., Yan, D., Chalouni, C., Luis, E., Khalfin, Y., Plise, E., Cheong, J., Lyssikatos, J. P., Strandh, M., Koefoed, K., Andersen, P. S., Flygare, J. A., Wah Tan, M., Brown, E. J., and Mariathasan, S. (2015) Novel antibody-antibiotic conjugate eliminates intracellular *S. aureus*. Nature. 527, 323–328

23. van Dalen, R., Molendijk, M. M., Ali, S., van Kessel, K. P. M., Aerts, P., van Strijp, J. A. G., de Haas, C. J. C., Codee, J., and van Sorge, N. M. (2019) Do not discard *Staphylococcus aureus* WTA as a vaccine antigen. Nature 572, E1–E2

24. Simanski, M., Glaser, R., Koten, B., Meyer-Hoffert, U., Wanner, S., Weidenmaier, C., Peschel, A., and Harder, J. (2013) *Staphylococcus aureus* subverts cutaneous defense by D-alanylation of teichoic acids. Exp Dermatol 22, 294–296

25. Bayer, A. S., Mishra, N. N., Cheung, A. L., Rubio, A., and Yang, S. J. (2016) Dysregulation of *mprF* and *dltABCD* expression among daptomycin-non-susceptible MRSA clinical isolates. J Antimicrob Chemother 71, 2100–2104

26. Ma, Z., Lasek-Nesselquist, E., Lu, J., Schneider, R., Shah, R., Oliva, G., Pata, J., McDonough, K., Pai, M. P., Rose, W. E., Sakoulas, G., and Malik, M. (2018) Characterization of genetic changes associated with daptomycin nonsusceptibility in *Staphylococcus aureus*. PLoS One 13, e0198366

27. Koprivnjak, T., Peschel, A., Gelb, M. H., Liang, N. S., and Weiss, J. P. (2002) Role of charge properties of bacterial envelope in bactericidal action of human group IIA phospholipase A2 against *Staphylococcus aureus*. J Biol Chem 277, 47636–47644

28. Winstel, V., Kuhner, P., Salomon, F., Larsen, J., Skov, R., Hoffmann, W., Peschel, A., and Weidenmaier, C. (2015) Wall Teichoic Acid Glycosylation Governs *Staphylococcus aureus* Nasal Colonization. mBio 6, 1–8

29. Brown, S., Xia, G., Luhachack, L. G., Campbell, J., Meredith, T. C., Chen, C., Winstel, V., Gekeler, C., Irazoqui, J. E., Peschel, A., and Walker, S. (2012) Methicillin resistance in *Staphylococcus aureus* requires glycosylated wall teichoic acids. Proc Natl Acad Sci U S A 109, 18909–18914

30. Xia, G., Maier, L., Sanchez-Carballo, P., Li, M., Otto, M., Holst, O., and Peschel, A. (2010) Glycosylation of wall teichoic acid in *Staphylococcus aureus* by TarM. J Biol Chem 285, 13405–13415

31. Gerlach, D., Guo, Y., De Castro, C., Kim, S. H., Schlatterer, K., Xu, F. F., Pereira, C., Seeberger, P. H., Ali, S., Codee, J., Sirisarn, W., Schulte, B., Wolz, C., Larsen, J., Molinaro, A., Lee, B. L., Xia, G., Stehle, T., and Peschel, A. (2018) Methicillin-resistant *Staphylococcus aureus* alters cell wall glycosylation to evade immunity. Nature. 563, 705–709

32. Xiong, M., Zhao, J., Huang, T., Wang, W., Wang, L., Zhao, Z., Li, X., Zhou, J., Xiao, X., Pan, Y., Lin, J., and Li, Y. (2020) Molecular Characteristics, Virulence Gene and Wall Teichoic Acid Glycosyltransferase Profiles of *Staphylococcus aureus:* A Multicenter Study in China. Front Microbiol 11, 2013

33. van der Es, D., Hogendorf, W. F., Overkleeft, H. S., van der Marel, G. A., and Codee, J. D. (2017) Teichoic acids: synthesis and applications. Chem Soc Rev 46, 1464–1482

34. Winstel, V., Liang, C., Sanchez-Carballo, P., Steglich, M., Munar, M., Broker, B. M., Penades, J. R., Nubel, U., Holst, O., Dandekar, T., Peschel, A., and Xia, G. (2013) Wall teichoic acid structure governs horizontal gene transfer between major bacterial pathogens. Nature communications 4, 2345

35. Wamhoff, E. C., Schulze, J., Bellmann, L., Rentzsch, M., Bachem, G., Fuchsberger, F. F., Rademacher, J., Hermann, M., Del Frari, B., van Dalen, R., Hartmann, D., van Sorge, N. M., Seitz, O., Stoitzner, P., and Rademacher, C. (2019) A Specific, Glycomimetic Langerin Ligand for Human Langerhans Cell Targeting. ACS Cent Sci 5, 808–820

36. Brown, E. J. (2015) Anti-wall teichoic antibodies and conjugates. Genentech, Inc., United States

37. Koc, C., Gerlach, D., Beck, S., Peschel, A., Xia, G., and Stehle, T. (2015) Structural and enzymatic analysis of TarM glycosyltransferase from *Staphylococcus aureus* reveals an oligomeric protein specific for the glycosylation of wall teichoic acid. J Biol Chem 290, 9874–9885

38. Masterson, A. J., Sombroek, C. C., De Gruijl, T. D., Graus, Y. M., van der Vliet, H. J., Lougheed, S. M., van den Eertwegh, A. J., Pinedo, H. M., and Scheper, R. J. (2002) MUTZ-3, a human cell line model for the cytokine-induced differentiation of dendritic cells from CD34+ precursors. Blood 100, 701–703

39. Santegoets, S. J., Masterson, A. J., van der Sluis, P. C., Lougheed, S. M., Fluitsma, D. M., van den Eertwegh, A. J., Pinedo, H. M., Scheper, R. J., and de Gruijl, T. D. (2006) A CD34(+) human cell line model of myeloid dendritic cell differentiation: evidence for a CD14(+)CD11b(+) Langerhans cell precursor. J Leukoc Biol 80, 1337–1344

40. Stemerding, A. M., Kohl, J., Pandey, M. K., Kuipers, A., Leusen, J. H., Boross, P., Nederend, M., Vidarsson, G., Weersink, A. Y., van de Winkel, J. G., van Kessel, K. P., and van Strijp, J. A. (2013) *Staphylococcus aureus* formyl peptide receptor-like 1 inhibitor (FLIPr) and its homologue FLIPr-like are potent FcgammaR antagonists that inhibit IgG-mediated effector functions. J Immunol 191, 353–362

41. van Dalen, R., Fuchsberger, F. F., Rademacher, C., van Strijp, J. A. G., and van Sorge, N. M. (2020) A Common Genetic Variation in Langerin (CD207) Compromises Cellular Uptake of *Staphylococcus aureus*. Journal of innate immunity 12, 191–200

42. Lacey, K. A., Mulcahy, M. E., Towell, A. M., Geoghegan, J. A., and McLoughlin, R. M. (2019) Clumping factor B is an important virulence factor during *Staphylococcus aureus* skin infection and a promising vaccine target. PLoS Pathog 15, e1007713

43. Hazenbos, W. L., Kajihara, K. K., Vandlen, R., Morisaki, J. H., Lehar, S. M., Kwakkenbos, M. J., Beaumont, T., Bakker, A. Q., Phung, Q., Swem, L. R., Ramakrishnan, S., Kim, J., Xu, M., Shah, I. M., Diep, B. A., Sai, T., Sebrell, A., Khalfin, Y., Oh, A., Koth, C., Lin, S. J., Lee, B. C., Strandh, M., Koefoed, K., Andersen, P. S., Spits, H., Brown, E. J., Tan, M. W., and Mariathasan, S. (2013) Novel staphylococcal glycosyltransferases SdgA and SdgB mediate immunogenicity and protection of virulence-associated cell wall proteins. PLoS Pathog 9, e1003653

44. Bleiziffer, I., Eikmeier, J., Pohlentz, G., McAulay, K., Xia, G., Hussain, M., Peschel, A., Foster, S., Peters, G., and Heilmann, C. (2017) The Plasmin-Sensitive Protein Pls in Methicillin-Resistant *Staphylococcus aureus* (MRSA) Is a Glycoprotein. PLoS Pathog 13, e1006110

45. Siboo, I. R., Chambers, H. F., and Sullam, P. M. (2005) Role of SraP, a Serine-Rich Surface Protein of *Staphylococcus aureus,* in binding to human platelets. Infect Immun 73, 2273–2280

46. Xia, G., Kohler, T., and Peschel, A. (2010) The wall teichoic acid and lipoteichoic acid polymers of *Staphylococcus aureus*. International Journal of Medical Microbiology 300, 148–154

47. Wanner, S., Schade, J., Keinhorster, D., Weller, N., George, S. E., Kull, L., Bauer, J., Grau, T., Winstel, V., Stoy, H., Kretschmer, D., Kolata, J., Wolz, C., Broker, B. M., and Weidenmaier, C. (2017) Wall teichoic acids mediate increased virulence in *Staphylococcus aureus*. Nat Microbiol 2, 16257

48. Mistretta, N., Brossaud, M., Telles, F., Sanchez, V., Talaga, P., and Rokbi, B. (2019) Glycosylation of *Staphylococcus aureus* cell wall teichoic acid is influenced by environmental conditions. Scientific reports 9, 3212

49. Li, X., Gerlach, D., Du, X., Larsen, J., Stegger, M., Kuhner, P., Peschel, A., Xia, G., and Winstel, V. (2015) An accessory wall teichoic acid glycosyltransferase protects *Staphylococcus aureus* from the lytic activity of Podoviridae. Scientific reports 5, 17219

50. McDermott, R., Bausinger, H., Fricker, D., Spehner, D., Proamer, F., Lipsker, D., Cazenave, J. P., Goud, B., De La Salle, H., Salamero, J., and Hanau, D. (2004) Reproduction of Langerin/CD207 traffic and Birbeck granule formation in a human cell line model. The Journal of investigative dermatology 123, 72–77

51. Pasparakis, M., Haase, I., and Nestle, F. O. (2014) Mechanisms regulating skin immunity and inflammation. Nat Rev Immunol 14, 289–301

52. Li, R. E., Hogervorst, T. P., Achilli, S., Bruijns, S. C., Arnoldus, T., Vives, C., Wong, C. C., Thepaut, M., Meeuwenoord, N. J., van den Elst, H., Overkleeft, H. S., van der Marel, G. A., Filippov, D. V., van Vliet, S. J., Fieschi, F., Codee, J. D. C., and van Kooyk, Y. (2019) Systematic Dual Targeting of Dendritic Cell C-Type Lectin Receptor DC-SIGN and TLR7 Using a Trifunctional Mannosylated Antigen. Front Chem 7, 650

