## Supplementary information for "The impact of *Staphylococcus aureus* cell wall glycosylation on langerin recognition and Langerhans cell activation"

**Supporting information**

List of materials included:

- Table S1
- Supporting figures 1-3

**Table S1 Bacterial strains used in this study**

| **Strain (sequence type, clonal complex)** | **Source** |
| --- | --- |
| N315 WT (ST5, CC5) | NARSA strain collection |
| N315 Δ*tarS* | (31) |
| N315 Δ*tarP* | (31) |
| N315 Δ*tarS*Δ*tarP* | (31) |
| N315 Δ*tarS*Δ*tarP+p*RB474*-tarS* | This study |
| N315 Δ*tarS*Δ*tarP+p*RB474*-tarP* | This study |
| RN4220 WT (ST8, CC8) | (53) |
| RN4220 Δ*dltA* | (54) |
| RN4220 Δ*tarM*Δ*tarS* | (34) |
| RN4220 Δ*tarM*Δ*tarS+*pRB474*-tarS* | (34) |
| RN4220 Δ*tarM*Δ*tarS+*pRB474*-tarP* | (31) |
| RN4220 Δ*tarM*Δ*tarS+*pRB474*-tarM* | (34) |

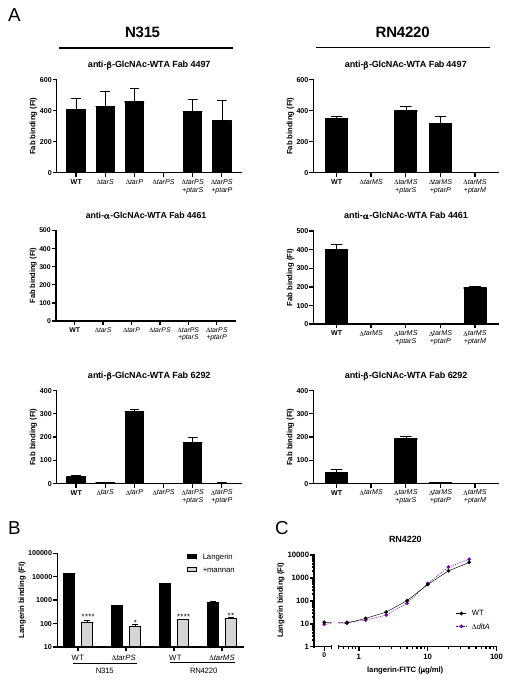

**Supporting Figure 1** A) Binding of Fab specific for α-GlcNAc-WTA (4461), β-GlcNAc-WTA (4497) and β-1,4-GlcNAc-WTA (6292) to N315 (left) and RN4220 (right) mutant panels. B) Binding of recombinant langerin-FITC (40 µg/ml) to N315 WT, N315 Δ*tarPS,* RN4220 WT and RN4220 Δ*tarMS* in the presence of absence of mannan (20 µg/ml). C) Binding of recombinant langerin-FITC (0.6-40 µg/ml) to RN4220 wt and Δ*dltA.* Data is depicted as geometric mean fluorescence intensity (MFI) + standard error of mean (SEM) of biological triplicates. **p* < 0.05, ***p* < 0.01, *****p* < 0.0001

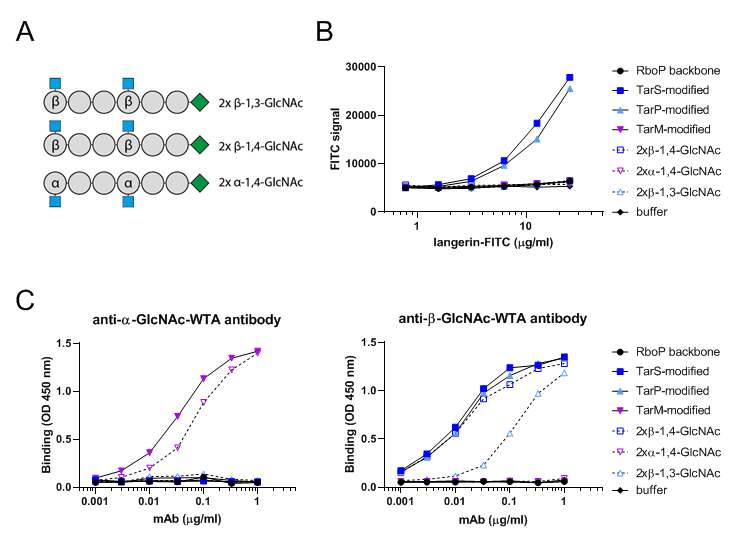

**Supporting figure 2** A) Schematic overview of fully synthetic WTA oligomers. Grey circles indicate RboP subunit, blue square represents GlcNAc, green diamond indicates biotin tag, α and β indicate the type of linkage of GlcNAc to RboP subunit. B) Binding of recombinant langerin-FITC (0.8-25 µg/ml) to *in vitro*-glycosylated RboP hexamers, fully synthetic WTA oligomers and RboP backbone. C) Binding of monoclonal antibodies (0.001-1 µg/ml) specific for α-GlcNAc-WTA (4461) and β-GlcNAc-WTA (4497) to the same panel of WTA oligomers as in B, followed by anti-human IgG-HRP staining. Data for panel B is shown as fluorescence signal, for C the absorbance at 450nm.

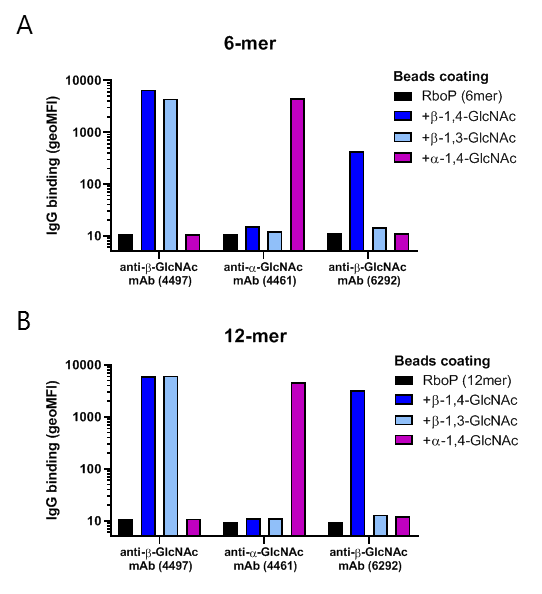

**Supporting figure 3** Binding of monoclonal antibodies (3 µg/ml) specific for α-GlcNAc-WTA (4461), β-GlcNAc-WTA (4497) and β-1,4-GlcNAc-WTA (6292) to beads coated with (A) RboP hexamers and (B) RboP dodecamers, *in vitro*-glycosylated by TarS, TarP or TarM. Unglycosylated RboP fragments are included as controls, binding is shown as geometric mean fluorescence intensity, after staining with goat anti-human kappa-Alexa Fluor 647.
